# Different Expressions of IL-1 TNF-α P-selectin mRNA by Endothelial Cells after Vein Thrombosis

**DOI:** 10.1101/681387

**Authors:** Jiasheng Xu, Kaili Liao, Weimin Zhou

## Abstract

**Objective:** Experiments were designed to compare the expressions of IL-1 TNF-α P-selectin mRNA by porcine endothelial cells after vein thrombosis.

**Methods:** IVC under the renal vein of 20 porcines were ligated to induce thrombosis modes. These thrombosed veins were divided into three groups:group A-one day after thrombosis, group B-four days after thrombosis and group C-seven days after thrombosis. The other pigs were given the shame operation as a contro group (group D). The mRNA levels of IL-1、 TNF-α and P-selectin expressed by porcine endothelial cells in three groups were analy sed by semi quantitative RT-PCR. Endothelial cells were harvested with collagenase II.

**Results:** The purity of endothelial cells harvested was 99.42 ±0.07. The expression of IL-1 was detained only in group A while TNF-αreached its peak in group B(*P* <0.05) and P-selectin increased gradually with the days passing by(*P* <0.05).

**Conclusion:** Endothelial cells are not only the target cells of inflammatory mediators, but also can express a variety of active factors to promote venous thrombosis. Expression of TNF-α mRNA is increased gradually in the early period of vein thrombosis while *P*-selectin in the acute period; IL-1 mRNA was transiently expressed only in the early stage of thrombosis.

## Introduction

With the improvement of people’s living standards and the improvement of examination methods, the incidence of deep vein thrombosis has also gradually increased, and it has become another common disease that endangers people’s health. It is well known that blood coagulation balance is maintained by the interaction between vascular endothelial cells, blood cells, and coagulation-fibrino systems, and cytokines and adhesion molecules are closely related to this interaction. In this experiment, SQ RT-PCR (semi-quantitative reverse transcription polymerase chain reaction) was used to detect IL-1 (interleukin-1), TNF-α (tumor necrosis factor-α) and P-selectin (P-Selectin, PS) mRNA expression changes at different time points of venous thrombosis to explore the mechanism of venous thrombosis.

## 1 Materials and methods

### 1.1 material

Take 20 piglets, male or female, weighing 20 ∼ 25kg. After 15 cases of the lower leg of the piglet (injected into the inferior vena cava plane), the inferior vena cava (the right renal vein was injected into the inferior vena cava plane to the inferior vena cava to the common iliac vein), a thrombus model was obtained. For group A (1d, n = 5 after thrombosis), group B (4d, n = 5 after thrombosis) and group C (7d after thrombus formation, n = 5), another 5 sham operation as control group D.

### 1.2 method

#### 1.2.1 Preparation of a thrombus model

Animals were placed in a supine position and fasted before surgery. After strict sterilization by aseptic technique, under the general anesthesia (intravenous injection of pentobarbital sodium 30mg kg), the inferior vena cava (IVC) was exposed through the lower abdomen longitudinal incision, and the segment was free of about 8cm, and the branches were ligated. The distal end of the the free inferior vena cava was ligated with a rubber band**(Figure 1)**. After 48 hours, the ligated rubber band was released again and the thrombus was confirmed by angiography[1]. Sham surgery means only revealing the kidney inferior segment of the inferior vena cava and ligating the branches of the genus but not ligating the proximal or distal end. Five piglets undergoing sham surgery were used as the control group.

**Figure 1.**
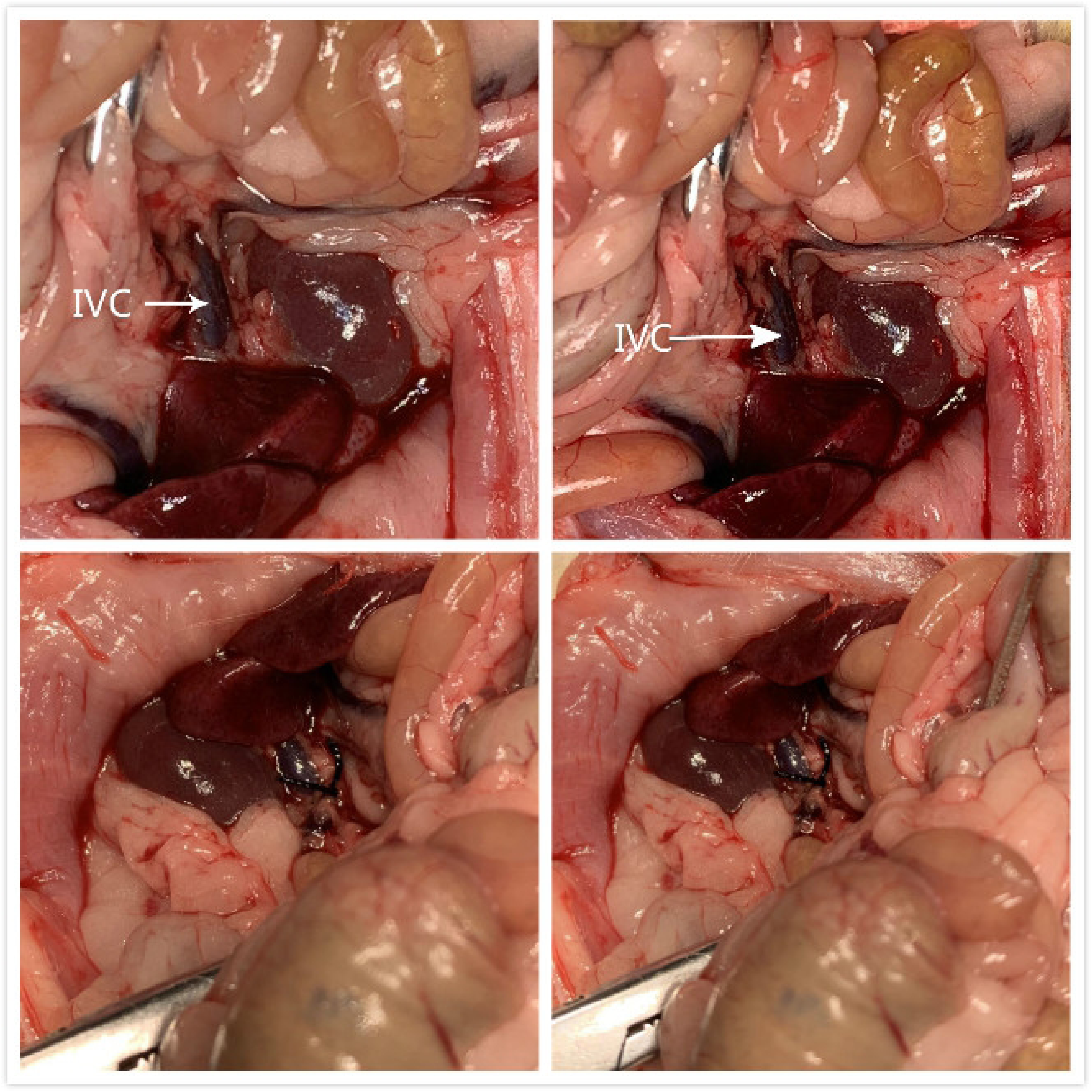
the inferior vena cava (IVC) was exposed through the lower abdomen longitudinal incision. The distal end of the the free inferior vena cava was ligated with a rubber band.

#### 1.2.2 Specimen collection

venography was performed on the 1st, 4th, and 7th day after porcine venous thrombosis, and thrombus formation was confirmed after thrombus formation. The scalp needle was inserted into the vascular lumen at both ends, and lava was lavaged three times to wash away residual blood cells. Then close one end and use a 10ml syringe to absorb 2 ∼ 3ml of type II collagenase pre-warmed at 37 °C. Inject 2μl of vascular RNA template 2μl, IL-1 or TNF-α or P-selectin and internal reference β-actin upstream and downstream primers (2μl (20 pmol), 1 μl of a mixture of TAQ reverse transcriptase and 18 μl of sterile double distilled water. The cycle conditions were 37 °C for 30 min, 94 °C for 2min for the first cycle, denaturation (94 °C, 30 s), annealing and closing both ends, and incubating in a 37 °C constant temperature water bath for 25 min[2-5]. After incubating, gently squeeze the blood vessel wall, collect the digestive juice in a 20 ml centrifuge tube, centrifuge at 1000 rpm for 10 min, discard the supernatant, and make the cell suspension with 10 ml of PBS solution.

#### 1.2.3 Counting and Purity Identification of Endothelial Cells

20 μl of cell suspension droplets were taken on a counting plate to calculate the total number of cells. Centrifuge the cell suspension at 0.5 ml for 10 min, fix 4 formaldehyde at 4 °C for 25 min, and wash the PBS solution three times. Rabbit anti-human VWF serum antibody was added, incubated at 37 °C for 30 min, and washed three times with PBS. Add goat anti-rabbit FITC-IgG secondary antibody, Incubate at 37 °C for 30 min and wash 3 times with PBS. Store in the dark, using flow cytometry within 2h. Each sample counted 10 000 cells from which approximately 8700 endothelial cells of a size and morphology within the normal range were analyzed. The computer collects the processed data. After the cell count and purity were identified, the cell suspension was centrifuged at 1000 rpm for 5 min, the supernatant was discarded, and the sinking cells were frozen at −80 °C in a centrifuge tube.

#### 1.2.4 Extraction of total RNA

1 ml of Trizol was added to a centrifuge tube in which a cell pellet was collected, repeatedly blown and vigorously mixed to dissolve the cells. Add 0.2 ml of chloroform, vigorously mix for 15 s, incubate at 15 ∼ 30 °C for 2 ∼ 3 min, centrifuge at 1200 r min for 15 min, and separate the mixture into the bottom red phenol-chloroform phase, mesophase and the upper layer of colorless aqueous phase containing total RNA. Transfer the aqueous phase to a new Eppendof tube, add 0.5 ml of isopropanol, incubate at 15 ∼ 30 °C for 10 min, and centrifuge for 10 min at 1200 rmin to reveal a jelly-like precipitate. The supernatant was discarded, washed with ethanol for 2 times, and the RNA was air-dried and dissolved in 100 μl of DEPC-pretreated double distilled water.

#### 1.2.5 SQ RT-PCR reaction and product electrophoresis

IL-1, TNF-α, P-selectin and β-actin (β actin) primers were synthesized according to references [6-9] (**Table 1**). According to the one-step PCR kit reaction instructions, the reaction volume contains 2 × RT-PCR Buffer 25μl RNA template 2μl, IL-1 or TNF-α or P-selectin and internal reference β-actin upstream and downstream primers total 2μl (20pmol 1 μl of a mixture of TAQ reverse transcriptase and 18 μl of sterile double distilled water. The cycle condition is 37 °C 30min, 94 °C 2min First cycle, denaturation (94 °C, 30s), annealing (58 °C, 30s), extension (68 °C, 1min) 35 cycles, 72 °C 6min last cycle. The above product was electrophoresed on a 1.5 agarose gel (90v, 1h), photographing under UV light, optical density scanning of semi-quantitative reaction products. The relative content of IL-1, TNF-α and P-selectin mRNA is expressed as the ratio of β-actin absorbance× area [10-13]

## 2 Results

2.1 The number of endothelial cells collected by enzymatic hydrolysis is about 10 μl, and the purity of endothelial cells is 99.42 ± 0.07 **(Figure. 2)**.

**Figure 2.**
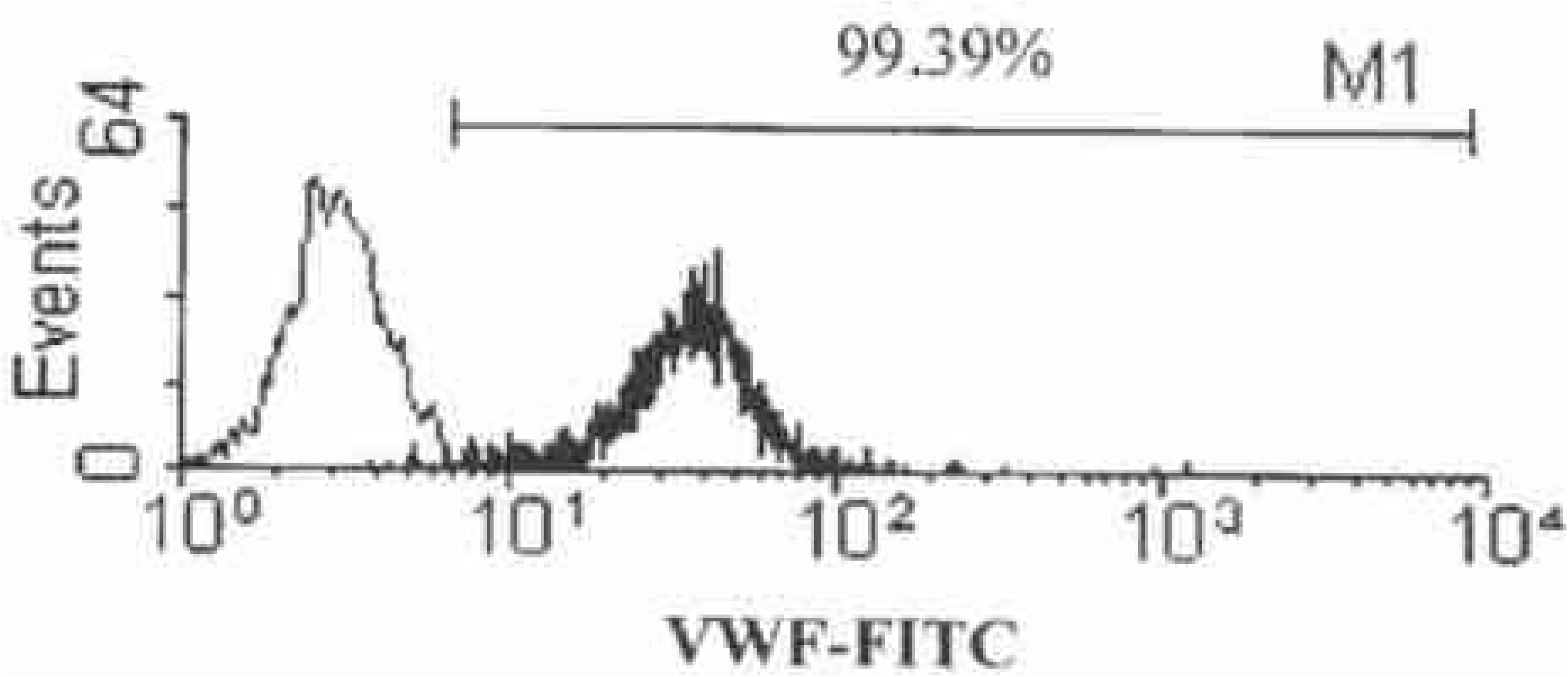
Flow cytometry to detect the purity of endothelial cells collected by enzymatic hydrolysis

2.2 SQ RT-PCR product electrophoresis results (**Figure 3, Table 2**)

**Figure 3.**
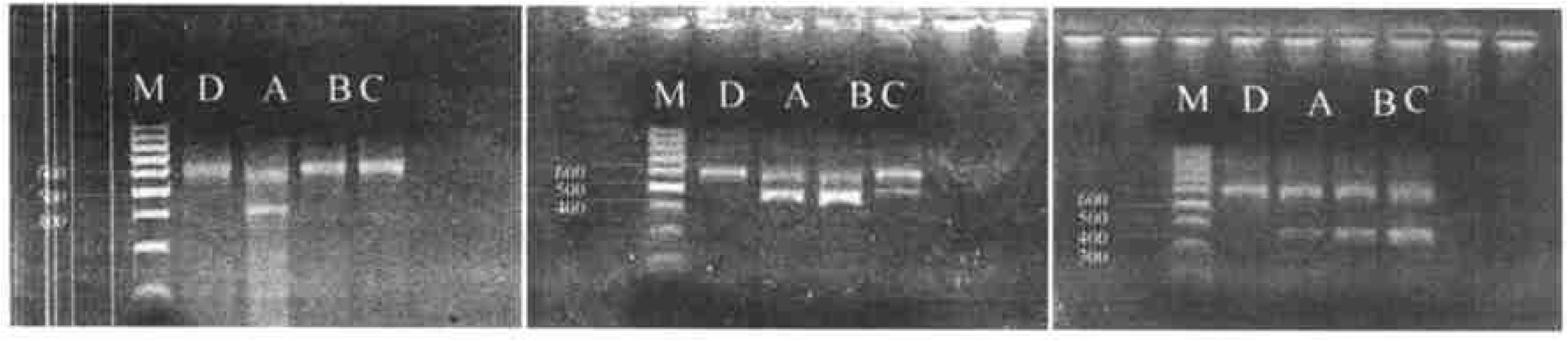
the left: RT-PCR detection of IL-1 mRNA expression; In the middle of the figure: RT-PCR detection of TNF-α mRNA expression; the right: RT-PCR detection of P-smRNA expression

(1) The control group only showed a band of β-actin mRNA. (2) IL-1 mRNA is only expressed in group A. (3) TNFα mRNA was expressed in group A, peaked in group B (the broadest and brightest of the product bands), and decreased in group C. (4) The expression of P-selectin mRNA gradually increased with the prolongation of time within 1 week of thrombosis. (5) Using an image analyzer to perform optical density scanning on the bands of β-actin in each group, the expression levels were consistent.

## 3 Discussion

In the past two decades, many useful studies have been done on the genetic etiology, imaging diagnosis and internal surgical treatment of venous thrombosis at home and abroad, but endothelial cells express cytokines and adhesion molecules at different time points during venous thrombosis[14-15]. The changes have not been reported in the literature. In this experiment, we used SQ RT-PCR method, and the expression level of β-actin was consistent after agarose gel electrophoresis, indicating that the reaction conditions of each group were basically the same and comparable. This study showed that three inflammatory factors were expressed in different degrees in each group, which proved that endothelial cells are not only the target cells of inflammatory mediators, but also can present antigens, secrete and express a variety of active factors, mediate cell adhesion, and promote Platelet activation and coagulation processes occur. In this experiment, the band at 434 bp of group A was bright and clear, indicating that IL-1 was involved in thrombosis after endothelial cells were stimulated by inflammation. Although there was no product band in group B and group C, the reaction conditions were the same, and the reaction was excluded. System problems. Kurt-Jone [16] and others believe that IL-1m-RNA is fast in transcription and translation. After 30 minutes of stimulation, intracellular expression can be detected. It peaks at 3 to 4 hours after stimulation, then decreases, and remains stable after 16 hours. Level, it will not be detected after 48h. This is consistent with the results of this experiment, and it is concluded that IL-1 plays a major role in the early stage of thrombosis[17-20]. The product band at 463 bp indicates that endothelial cells began to express TNF-α in the early stage of thrombus formation; the bands in the same position in group B were brighter and wider than those in group A, and there was a significant difference in optical density scanning analysis, indicating that In the prolongation of the course of thrombosis, TNF-α only acts on the early stage of venous thrombosis[21-23], and on the other hand may be related to the commonality of most cytokines, that is, they are generally detected after 6-8 hours of activation, 24 ∼ 72h. After reaching the peak, then the reduction is maintained at a certain level. P-selectin is currently the most specific marker for reflecting platelet activation and release, and plays a central role in inflammation and embolism [24-25]. In this experiment, the band of 342 bp in group A agarose gel electrophoresis indicates that P-selectin is expressed by endothelial cells in the early stage of thrombosis, and participates in the activation and adhesion of platelets and leukocytes. The product bands in the same position in B and C showed progressive brightening and broadening, indicating that P-selectin is continuously expressed and gradually enhanced during thrombus formation, and is one of the important inflammatory factors that promote thrombosis and development. The results of the D group showed that the expression of TNF-α, IL-1 and P-selectin mRNA was that the vascular endothelial cells were first activated.

## 4 Conclusions

Through this experiment, the following conclusions can be drawn: (1) Endothelial cells are not only the target cells of inflammatory mediators, but also can express a variety of active factors to promote venous thrombosis. (2) IL-1 mRNA was transiently expressed only in the early stage of thrombosis; TNF-α mRNA expression gradually increased in the early stage of thrombosis; P-selectin mRNA expression gradually increased in the acute phase of thrombosis.

## Declarations

### Availability of data and supporting materials

Please contact author for data requests.

### Ethical Approval and Informed consent

#### Ethical approval

This article does not contain any studies with human performed by any of the authors. All procedures performed in studies involving animals participants were in accordance with the ethical standards of the institutional and/or national research committee. The study was approved by the ethics committee of the Second Affiliated Hospital of Nanchang University.

### Consent for publication

Not applicable.

### Author contribution statement

Author:Jiasheng Xu Contribution:

1. Design research direction
2. Writing papers

Author:KAILI LIAO Contribution:

1. searching for references
2. helping to write papers

Author:WEIMIN ZHOU Contribution:

1. Review and revise the papers
2. Guidance article writing

## Acknowledgements

None

